# Inactivating conditions of therapeutic mycobacteriophages

**DOI:** 10.1101/2025.09.30.679648

**Authors:** Andrew Wiggins, Umar N. Chaudhry, Fabiana Bisaro, Addison Lueck, Alan A. Schmalstig, Graham F. Hatfull, David B. Hill, Miriam Braunstein

**Affiliations:** Mycobacteria Research Laboratories, Department of Microbiology, Immunology, and Pathology, Colorado State University, Fort Collins, Colorado, USA; Department of Microbiology and Immunology, University of North Carolina at Chapel Hill, Chapel Hill, North Carolina, USA; Cell and Molecular Biology Program, Colorado State University, Fort Collins, Colorado, USA; Department of Biological Sciences, University of Pittsburgh, Pittsburgh, PA, USA; Marsico Lung Institute and Lampe Joint Department of Biomedical Engineering, University of North Carolina at Chapel Hill, Chapel Hill, North Carolina, USA

**Author notes:** Co-first authors (Andrew Wiggins, Umar N. Chaudhry).

**Keywords:** Bacteriophage, Nontuberculous Mycobacteria, mycobacteriophage

## Abstract

There is a need for new therapies to treat drug resistant nontuberculous mycobacteria (NTM) disease. Bacteriophages (phages), which are viruses that infect and kill bacteria, are actively being explored as an alternative approach for treating mycobacterial diseases. Several compassionate-use cases of phage therapy for drug resistant NTM infections exhibit favorable outcomes. To further the development of phage therapy it is important to recognize and avoid conditions that negatively impact phage activity during phage production, storage, formulation, or treatment. Conversely, there is a need to inactivate free phages in certain preclinical phage therapy experiments. In this study, we investigated three mycobacteriophages BPsΔ*33*HTH-HRM10, Muddy, and ZoeJΔ*45* from compassionate-use NTM treatment cases for their sensitivity to a variety of conditions that included temperature, acid pH, detergents, mucus, and phage inactivating buffers. Several conditions resulted in dramatic and rapid reductions in the level of active phage while others had no effect. We also observed different sensitivities between the phages. The results provide valuable information to support further investigation and development of these phages as therapeutics.

## Importance

Bacteriophages (phages) offer a promising alternative therapy for treating drug resistant mycobacterial infections. For successful implementation of phage therapy, it is important to recognize conditions that inactivate the phages. Here, we studied three mycobacteriophages from recent compassionate-use phage therapy cases for their sensitivity to a range of conditions that may be encountered in production, storage, formulation, or treatment. The results demonstrate sensitivity to some conditions and tolerance to others, and they additionally reveal phage-specific differences in sensitivities highlighting the need for direct evaluation of individual therapeutic phages during development.

## Observation

As bacteria get harder to treat with antibiotics, the need for alternative therapies becomes more urgent. An alternative to antibiotics is the use of lytic bacteriophages (phages), which infect and kill bacteria. There are a growing number of compassionate-use cases of phages being used therapeutically to treat patients with multidrug-resistant bacterial infections, including nontuberculous mycobacteria (NTM) pulmonary infections (1, 2) For phage therapy to succeed, it is important to avoid conditions that negatively impact phage activity when producing, storing, or formulating phages since low phage titers could jeopardize treatment, as likely occurred in a clinical trial of phages for *Pseudomonas aeruginosa* infections (3-5). Conversely, a method for inactivating phages is useful in preclinical experiments where the bactericidal activity of phages is measured by bacterial CFU (i.e., to avoid free phages from reducing CFU during outgrowth). Here, we evaluated three siphovirus mycobacteriophages from phage therapy cases of NTM infections, each mapping to a different genomic cluster, for their sensitivity to a variety of conditions: BPsΔ*33*HTH-HRM10 (BPsΔ) (cluster G), Muddy (cluster AB), and ZoeJΔ*45* (ZoeJΔ) (cluster K) (6).

To measure phage sensitivities, phages at titers of 10^9^ – 10^10^ PFU/ml in mycobacteriophage (MP) buffer were exposed to different conditions at room temperature (22°C), unless otherwise indicated. At specific time points, phages were quantified as PFU/ml using an agar overlay assay with *Mycobacterium smegmatis*. Conditions resulting in a significant reduction in titer compared to an untreated phage control at the same time point were determined by calculating the log change (log PFU_challenge_ – log PFU_control_) (see Supplemental File for Materials and Methods). All data from replicate experiments are presented in Table 1.

**Table 1:** Mycobacteriophage Sensitivities (Log PFU/ml_Treatment_ - Log PFU/ml_Control_)

|  |  |  |  |  |
| --- | --- | --- | --- | --- |
| **Condition** | **Time** | **BPsΔ** | **Muddy** | **ZoeJΔ** |
| 37°C | 1, 5, 10, 60, 120 min | (NS, NS) | (NS, NS) | (NS, NS) |
| 45°C | 1,5,10,60, 120 min | (NS, NS) | (NS, NS) | (NS, NS) |
| 55°C | 1 min | (NS, NS) | (NS, NS) | (NS, NS) |
| 55°C | 5 min | **(-1.09, -1.27)** | (NS, NS) | (NS, NS) |
| 55°C | 10 min | **(-1.47, -1.47)** | (NS, NS) | (NS, NS) |
| 55°C | 60 min | **(-2.64, -2.59)** | (NS, NS) | (NS, NS) |
| 55°C | 120 min | **(-2.97, -3.12)** | **(-1.58, -2.11)** | **(-1.04, -1.16)** |
| 70°C | 1 min | **(-5.40, -6.74)** | **(-1.90, -1.03)** | **(-4.13, -4.99)** |
| 70°C | 5 min | **(BLD, BLD)** | **(-5.84, -5.72)** | **(-4.90, -5.13)** |
| 70°C | 10 min | **(BLD, BLD)** | **(BLD, BLD)** | **(-4.78, BLD)** |
| 70°C | 60 min | **(BLD, BLD)** | **(BLD, BLD)** | **(BLD, BLD)** |
| 70°C | 120 min | **(BLD, BLD)** | **(BLD, BLD)** | **(BLD, BLD)** |
| 90%EtOH | 1, 5, 10, 60min | **(BLD, BLD)** | **(BLD, BLD)** | **(BLD, BLD)** |
| 63% EtOH | 1 min | **(-5.56, -5.13)** | **(-4.36, -2.16)** | **(-2.93, -2.42)** |
| 63% EtOH | 5 min | **(-6.47, -6.14)** | **(-4.72, -3.75)** | **(-3.80, -2.68)** |
| 63% EtOH | 10 min | **(BLD, -6.28)** | **(-4.36, -4.28)** | **(-3.14, -3.28)** |
| 63% EtOH | 60 min | **(BLD, BLD)** | **(-3.97, -4.79)** | **(-4.74, -5.19)** |
| 1:10 PIB pH 3 | 1 min | **(BLD, BLD)** | **(BLD, BLD)** | **(-6.19, -4.55)** |
| 1:10 PIB pH 3 | 5 min | **(BLD, BLD)** | **(BLD, BLD)** | **(BLD, BLD)** |
| 1:10 PIB pH 3 | 10 min | **(BLD, BLD)** | **(BLD, BLD)** | **(BLD, BLD)** |
| 1:10 PIB pH 3 | 60 min | **(BLD, BLD)** | **(BLD, BLD)** | **(BLD, BLD)** |
| 1:10 PIB pH 4 | 1 min | **(-1.71, -1.86)** | (NS, NS) | (NS, NS) |
| 1:10 PIB pH 4 | 5 min | **(-2.02, -2.08)** | (NS, NS) | (NS, NS) |
| 1:10 PIB pH 4 | 10 min | **(-2.43, -2.30)** | (NS, NS) | (NS, NS) |
| 1:10 PIB pH 4 | 60 min | **(-3.78, -3.67)** | **(-0.53, NS)** | (NS, NS) |
| 1:10 PIB pH 5 | 1 min | (NS, NS, NS) | (NS, NS) | (NS, NS) |
| 1:10 PIB pH 5 | 5 min | (NS, NS, NS) | (NS, NS) | (NS, NS) |
| 1:10 PIB pH 5 | 10 min | (NS, NS, NS) | (NS, NS) | (NS, NS) |
| 1:10 PIB pH 5 | 60 min | **(-0.62, -0.66, -0.67)** | (NS, NS) | (NS, NS) |
| MP pH 2 | 1, 5, 10, 60 min | **(BLD, BLD)** | **(BLD, BLD)** | **(BLD, BLD)** |
| MP pH 2 | 24 hrs | **(BLD, BLD)** | **(BLD, BLD)** | **(BLD, BLD)** |
| MP pH 3 | 1, 5, 10, 60 min | (NS, NS) | (NS, NS, NS) | (NS, NS) |
| MP pH 4 | 1, 5, 10, 60 min | (NS, NS) | (NS, NS) | (NS, NS) |
| MP pH 5 | 1, 5,10, 60 min | (NS, NS) | (NS, NS) | (NS, NS) |
| 2.25 mM FAS+0.002% TA | 1 min | **(BLD, BLD)** | **(BLD, BLD)** | **(BLD, BLD)** |
| 2.25 mM FAS | 1 min | **(-4.26, -4.55)** | **(BLD, BLD)** | **(-5.18, -5.23)** |
| 0.002% TA | 3 min | (NS, NS) | (NS, NS) | (NS, NS) |
| 2% HBE Mucus | 24 hrs | (NS, **-0.40**, NS) | (NS, NS) | (NS, NS) |
| 4% HBE Mucus | 24 hrs | **(-0.15, -0.32, NS)** | (NS, NS) | (NS, NS) |
| 6% HBE Mucus | 24 hrs | **(-0.40, -0.48, -0.69)** | (NS, **-0.60**) | (NS, NS) |
| 10 mM TCEP pH 1.9 | 24 hrs | **(BLD, BLD)** | **(BLD, BLD)** | **(BLD, BLD)** |
| 10 mM TCEP pH 7 | 24 hrs | (NS, NS) | (NS, NS) | (NS, NS) |
| 1 mM DTT pH 5.3 | 24 hrs | **(-5.2, -5.0, -5.2, -5.0)** | **(-0.32, -0.56)** | **(-3.67, -3.41)** |
| 1 mM DTT pH 7.0 | 24 hrs | (NS, NS) | (NS, NS) | **(-0.45, -2.64)** |
| 0.1% Tween 80 | 1 min | (NS, NS) | (NS, NS) | (NS, NS) |
| 0.1% Tween 80 | 5 min | (NS, NS) | (NS, NS) | (NS, NS) |
| 0.1% Tween 80 | 10 min | (NS, NS) | (NS, NS) | (NS, NS) |
| 0.1% Tween 80 | 60 min | (NS, NS) | (NS, NS) | (NS, **-0.57**) |
| 0.1% Tyloxapol | 1, 5, 10, 60 min | (NS, NS) | (NS, NS) | (NS, NS) |
| 0.1% Triton X-100 | 1 min | (NS, NS, NS, NS) | (NS, NS, NS, NS) | (NS, NS, NS, NS) |
| 0.1% Triton X-100 | 5 min | (NS, NS, NS, NS) | (NS, NS, NS, NS) | (NS, NS, NS, NS) |
| 0.1% Triton X-100 | 10 min | (NS, NS, NS, NS) | (NS, NS, NS, NS) | (NS, NS, NS, NS) |
| 0.1% Triton X-100 | 60 min | (NS, **-1.46**, NS, NS) | (NS, NS, NS, NS) | (NS, NS, NS, NS) |
| 1% Triton X-100 | 1 min | (NS, NS) | (NS, NS) | **(-0.52, NS)** |
| 1% Triton X-100 | 5 min | **(+0.45, NS)** | (NS, NS) | (NS, NS) |
| 1% Triton X-100 | 10 min | (NS, NS) | (NS, NS) | (NS, NS) |
| 1% Triton X-100 | 60 min | (NS, NS) | (NS, NS) | (NS, **-0.67**) |
| 10% Triton X-100 | 1, 5, 10, 60 min | (NS, NS) | (NS, NS) | (NS, NS) |
| 1% DMSO | 1 min | (NS, NS) | (NS, NS) | (NS, NS) |
| 1% DMSO | 5 min | (NS, NS) | (NS, NS) | (NS, NS) |
| 1% DMSO | 10 min | (NS, NS) | (NS, **-0.56**) | (NS, NS) |
| 1% DMSO | 60 min | (NS, NS) | (NS, NS) | (NS, NS) |
| 1% DMSO | 24 hrs | (NS, NS) | (NS, NS) | (NS, NS) |
| 1% DMSO | 48 hrs | (NS, NS) | (NS, NS) | (NS, NS) |
| 1% DMSO | 72 hrs | (NS, NS) | (NS, NS) | (NS, NS) |

Bold text indicates significant difference compared to control based on (i) one-way ANOVA and Tukey's post hoc ( $p < 0.05$ ) or (ii) Unpaired t-test for conditions with a single time point ( $p < 0.05$ ). NS = No significant difference. BLD = Below limit of detection [2,500 = PFU/ml; Log (PFU/ml) =3.5].

## Temperature

Sensitivity to elevated temperature was monitored over a 120-minute (min) time course and compared to room temperature (22ºC) (Fig 1A). At 37ºC and 45ºC there was no significant reduction in titer for any of the phages. However, with 55ºC there was a time dependent reduction in all the phages, with BPsΔ exhibiting the greatest sensitivity. All three phages were highly sensitivity to 70ºC.

**Figure 1.**
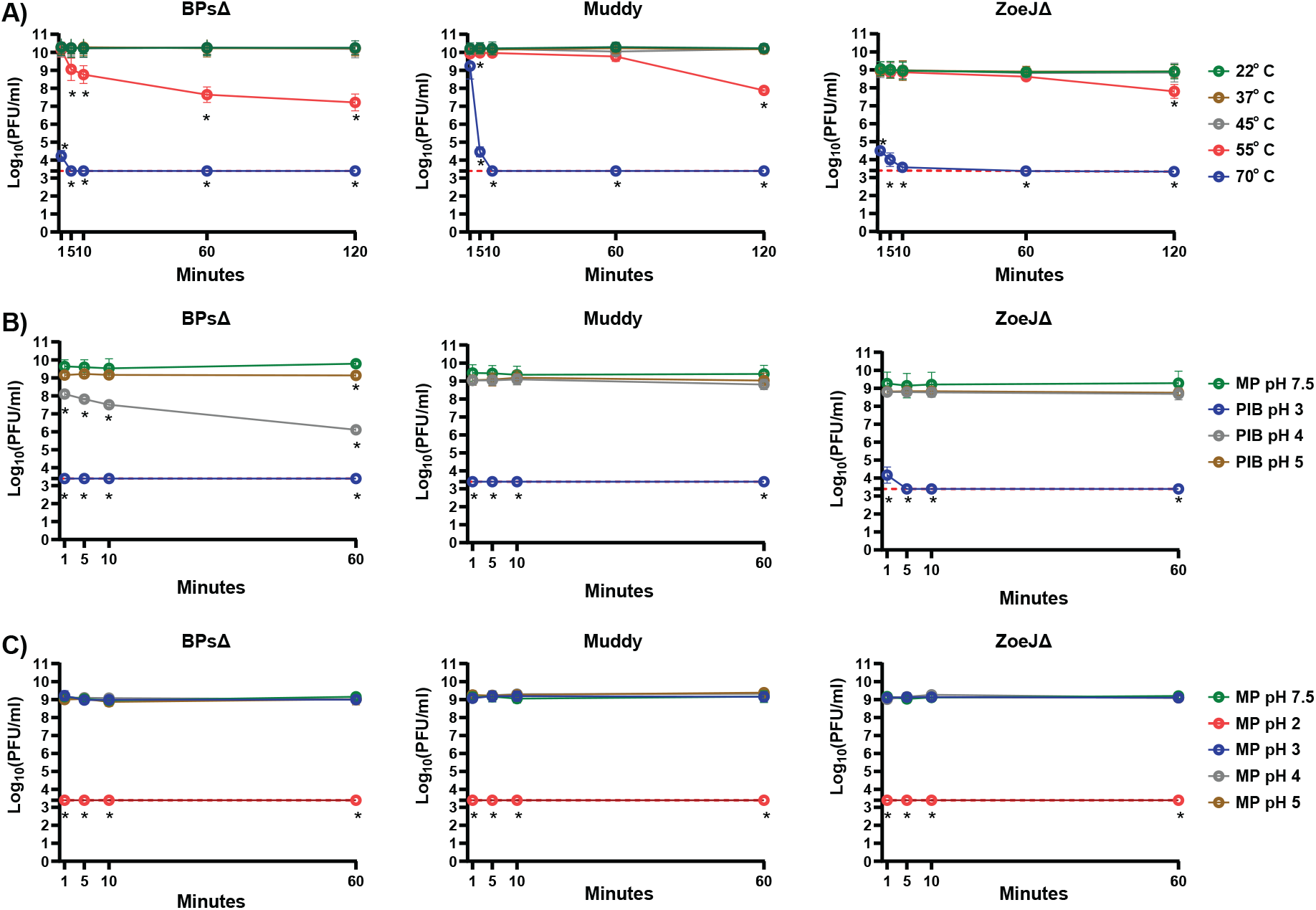
**A. Sensitivity of phages to various temperatures.** BPsΔ, Muddy, and ZoeJΔ were treated with a range of temperatures from to 22° C – 70° C. At specific time points, samples were removed and phages quantified. The mean of two independent experiments, each with three technical replicates, is plotted. Asterisks indicate significant reduction in log (PFU/ml) between a given treatment compared to the 22° C control at the same time point determined by one-way ANOVA (p<0.05) with Tukey’s post-test. **B. Effectiveness of Phage Inhibition Buffer (PIB) at various pH**. BPsΔ, Muddy, and ZoeJΔ were treated at a 1:10 ratio with PIB at pH 3, 4, and 5 and compared to MP Buffer pH 7.5. At specific time points, phages were quantified. The mean of two to three independent experiments, each with three technical replicates, is plotted. Asterisks indicate significant reduction in log (PFU/ml) between a given PIB pH time point compared to the MP buffer pH 7.5 control at the same time point determined by one-way ANOVA (p<0.05) with Tukey’s post-test. **C. Sensitivity of phages to various pH in MP buffer**. BPsΔ, Muddy, and ZoeJΔ were incubated in MP buffer at pH *2*, 3, 4, and 5 and compared to MP Buffer pH 7.5. At specific time points, samples were removed and phage quantified. The mean of two to three independent experiments, each with three technical replicates, was plotted. Asterisks indicate a significant reduction in log (PFU/ml) between a given pH time point compared to the MP buffer pH 7.5 control at the same time point determined by one-way ANOVA (p < 0.05) with Tukey’s post-test. The red dashed line indicates the limit of detection. Error bars indicate standard deviation. Samples with zero PFU recovered are plotted at the level of detection.

## Ethanol

Sensitivity to 63% and 90% ethanol, a commonly used disinfectant, was tested (Table 1). All three phages were highly sensitive to 90% ethanol with undetectable levels at 1 min. 63% ethanol also had a rapid, although incomplete, effect with BPsΔ again exhibiting the greatest sensitivity.

## Phage inactivation buffer (PIB) and acidic pH

When measuring phage bactericidal activity by reductions in CFU it is advantageous to inactivate free phage before CFU plating. We previously reported use of a 10 minute treatment with a Phage Inactivating Buffer (PIB) (940 mM citric acid, 10 mM potassium chloride, 135 mM NaCl, pH 3.0) to inactivate BPsΔ, Muddy, and ZoeJΔ (7-9). Importantly, the host bacteria of these phages *Mycobacterium abscessus* and *M. smegmatis* are not inhibited by PIB pH 3, 10 min (8) (Supplemental Fig 1). To determine the time-dependence of inactivation, we performed a PIB time course and observed rapid reduction of the phages to undetectable levels: by 1 min for BPsΔ and Muddy and 5 min for ZoeJΔ (Fig 1B). The effect of pH could be important (10, 11). Therefore, we investigated the adjustment of PIB to either pH 4 or pH 5. Compared to PIB pH 3, PIB pH 4 and 5 had no effect on Muddy or ZoeJΔ, and PIB pH 4 and 5 was less effective at reducing BPsΔ (Fig. 1B). BPsΔ was again more sensitive than the other phages. We also studied the impact of acidic pH in MP buffer. All the phages were highly sensitive to MP buffer pH 2 (i.e., undetectable at 1 min) and resistant to MP pH 3, 4, and 5 (Fig. 1C). Thus, PIB pH 3 fully inactivates but MP pH 3 does not, which indicates that the pH of PIB is important but not the sole factor responsible for phage inhibition.

## Ferrous ammonium sulfate (FAS) +/-tannic acid (TA)

FAS +/-TA is another reported phage inactivating treatment (12-14). After 1 min incubation in 2.25 mM FAS with 0.002% TA the titer of all three phages was below the level of detection (Table 1). One min treatment with 2.25 mM FAS lacking TA also significantly reduced phage titers with Muddy being undetectable and BPsΔ and ZoeJΔ exhibiting a 4-5 log reduction. Treatment with 0.002% TA for 3 min had no effect (Table 1).

## Mucus

Several patients receiving phage therapy for pulmonary NTM disease have muco-obstructive lung diseases (2). Studies of the effect of mucus on other phages reveal inhibitory or beneficial effects (15-17). This led us to test the effect of mucus harvested from cultured human bronchial epithelial (HBE) cells (18, 19). Phages were incubated for 24 hours with 2%, 4%, or 6% mucus, which are concentrations that model healthy, moderate, and severe airway disease mucus (20, 21). Across the concentrations, all three phages exhibited either modest (≤0.7 log) or no significant reduction in titer. This finding of little to no effect of mucus supports the therapeutic use of these phages in people with muco-obstructive diseases.

## Reducing agents

Reducing agents that disrupt disulfide bonds degrade the mucin network of mucus. There is interest in developing reducing agents into mucolytics for muco-obstructive diseases (21, 22). At concentrations that reduce the viscosity of mucus, we tested reducing agents Tris(2-carboxyethyl)phosphine (TCEP) and Dithiothreitol (DTT) on phage activity (23). Because TCEP and DTT solutions are acidic, we tested each agent as an unbuffered solution and at pH 7. Without buffering, both 10 mM TCEP pH 1.9 and 1 mM DTT pH 5.3 after 24 hours significantly reduced the level of all three phages. However, both agents at pH 7 had no effect, apart from a modest reduction in ZoeJΔ PFU with 1mM DTT (Table1). Because mucus has a high buffering capacity, which limits pH effects *in vivo* (21), these results suggest that concurrent treatment with reducing mucolytics will not inhibit these phages.

## Detergents

Sensitivity to detergents encountered in mycobacterial experiments (Tween-80, Tyloxapol, Triton X-100) was investigated. Tween-80 and Tyloxapol are included in mycobacterial liquid media to promote dispersed growth. Triton X-100 is used to lyse cultured eukaryotic cells infected with mycobacteria (8). All three phages were resistant to 60 min incubation with 0.1% Tween-80, 0.1% Tyloxapol, or 0.1%, 1%, or 10% Triton X-100 (Table 1). This resistance was surprising since Tween-80 is reported to prevent mycobacteriophage D29 infection (24). We wondered if the detergents might not inhibit the phage directly but, rather, prevent phage infection through effects on the mycobacterial host. To test this idea, immediately prior to use, 0.1% Tween-80, 0.1% Tyloxapol, or no detergent was added to the *M. smegmatis* culture used in the overlay for phage quantification (Fig 2). For BPsΔ and ZoeJΔ, Tween-80 or Tyloxapol in the culture did not affect PFU number although Tyloxapol delayed the time for ZoeJΔ plaques to appear. In contrast, inclusion of Tween-80 or Tyloxapol in the culture inhibited Muddy PFU. These results demonstrate that the activity of some, but not all, phages are impacted by detergent-specific effects on mycobacteria.

**Figure 2.**
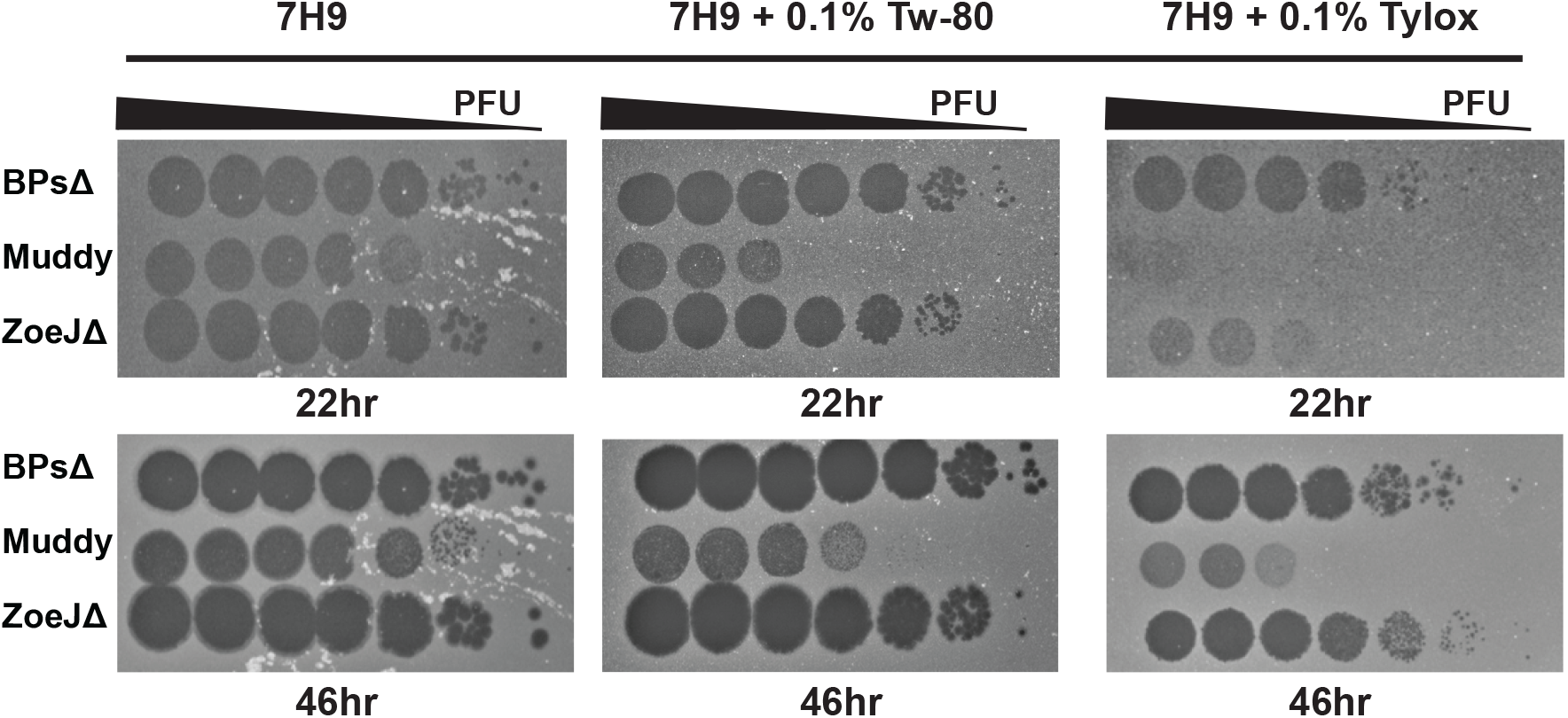
Sensitivity of phages to inclusion of detergent in the *M. smegmatis* culture used in agar overlays. BPsΔ, Muddy, and ZoeJΔ 10-fold serial dilutions spotted onto agar overlays prepared with *M. smegmatis* resuspended in Middlebrook 7H9 media supplemented with 0.5% glycerol, 0.2% glucose (no detergent) or the same Middlebrook 7H9 media with freshly added 0.1% Tween-80 (Tw-80) or 0.1% Tyloxapol (Tylox). Plates were incubated at 37° C for 22 and 46 hours. A representative image from two experiments is shown. All images were taken with the same exposure time of 0.09 seconds using a Bio-RAD GelDOC Go.

## DMSO

Many antimycobacterial drugs are dissolved in DMSO, such that 1% DMSO is likely to be included when phage-antibiotic combinations are tested on mycobacteria. DMSO may also be used as a protectant in long-term phage storage (25). Up to 72 hours, all three phages were resistant to 1% dimethyl sulfoxide (DMSO) (Table 1).

In conclusion, in studying three mycobacteriophages from phage therapy cases (2) we discovered several conditions that result in dramatic and rapid reductions in phage activity. We also observed differences between phages emphasizing the need for such studies. The results provide valuable information to support further evaluation of these phages as therapeutics for NTM disease.

## DATA AVAILABILITY

Data from this study will be made available through Dryad upon acceptance.

## Ethics Statement

HBE cell cultures are considered non-human subjects under the University of North Carolina Office of Research Ethics Biomedical Institutional Review Board protocol 03-1396, and the mucus harvested from them that was used in this project is likewise not considered mucus specimen under the University of North Carolina Office of Research Ethics Biomedical Institutional Review Board protocol 22-0990.

## Acknowledgments

M.B. acknowledges support from NIAID R01 AI176834, NIAID R21AI163677, and Cystic Fibrosis Foundation grant BRAUNS21P0. U.C. acknowledges support from NSF REU 2048087 and A.A.S. acknowledges support from a Kirschstein National Research Service Award (T32AI007151). D.B.H acknowledges support from the Cystic Fibrosis Foundation grant ESTHER24R0 and HILL25G0 and from NIH P30DK065988 We thank members of the Braunstein Lab for reviewing the manuscript. GFH was supported by grants from NIH (GM116884), Howard Hughes Medical Institute (GT12053), Cystic Fibrosis Foundation (Hatfull19GO and Hatfull21GO), and Emily’s Entourage. The authors declare no conflict of interest.

## References

1. Strathdee SA, Hatfull GF, Mutalik VK, Schooley RT. 2023. Phage therapy: From biological mechanisms to future directions. Cell 186:17–31.

2. Dedrick RM, Smith BE, Cristinziano M, Freeman KG, Jacobs-Sera D, Belessis Y, Whitney Brown A, Cohen KA, Davidson RM, van Duin D, Gainey A, Garcia CB, Robert George CR, Haidar G, Ip W, Iredell J, Khatami A, Little JS, Malmivaara K, McMullan BJ, Michalik DE, Moscatelli A, Nick JA, Tupayachi Ortiz MG, Polenakovik HM, Robinson PD, Skurnik M, Solomon DA, Soothill J, Spencer H, Wark P, Worth A, Schooley RT, Benson CA, Hatfull GF. 2022. Phage Therapy of Mycobacterium Infections: Compassionate-use of Phages in Twenty Patients with Drug-Resistant Mycobacterial Disease. Clin Infect Dis doi:10.1093/cid/ciac453.

3. Blazanin M, Lam WT, Vasen E, Chan BK, Turner PE. 2022. Decay and damage of therapeutic phage OMKO1 by environmental stressors. PLoS One 17:e0263887.

4. Jonczyk-Matysiak E, Lodej N, Kula D, Owczarek B, Orwat F, Miedzybrodzki R, Neuberg J, Baginska N, Weber-Dabrowska B, Gorski A. 2019. Factors determining phage stability/activity: challenges in practical phage application. Expert Rev Anti Infect Ther 17:583–606.

5. Jault P, Leclerc T, Jennes S, Pirnay JP, Que YA, Resch G, Rousseau AF, Ravat F, Carsin H, Le Floch R, Schaal JV, Soler C, Fevre C, Arnaud I, Bretaudeau L, Gabard J. 2019. Efficacy and tolerability of a cocktail of bacteriophages to treat burn wounds infected by Pseudomonas aeruginosa (PhagoBurn): a randomised, controlled, double-blind phase 1/2 trial. Lancet Infect Dis 19:35–45.

6. Dedrick RM, Guerrero-Bustamante CA, Garlena RA, Russell DA, Ford K, Harris K, Gilmour KC, Soothill J, Jacobs-Sera D, Schooley RT, Hatfull GF, Spencer H. 2019. Engineered bacteriophages for treatment of a patient with a disseminated drug-resistant Mycobacterium abscessus. Nat Med 25:730–733.

7. Brindley MA, Maury W. 2008. Equine infectious anemia virus entry occurs through clathrin-mediated endocytosis. J Virol 82:1628–37.

8. Schmalstig AA, Wiggins A, Badillo D, Wetzel KS, Hatfull GF, Braunstein M. 2024. Bacteriophage infection and killing of intracellular Mycobacterium abscessus. mBio 15:e0292423.

9. Zhang L, Sun L, Wei R, Gao Q, He T, Xu C, Liu X, Wang R. 2017. Intracellular Staphylococcus aureus Control by Virulent Bacteriophages within MAC-T Bovine Mammary Epithelial Cells. Antimicrob Agents Chemother 61.

10. Jeon G, Ahn J. 2021. Evaluation of phage adsorption to Salmonella Typhimurium exposed to different levels of pH and antibiotic. Microb Pathog 150:104726.

11. Dabrowska K. 2019. Phage therapy: What factors shape phage pharmacokinetics and bioavailability? Systematic and critical review. Medicinal Research Reviews 39:2000–2025.

12. McNerney R, Wilson SM, Sidhu AM, Harley VS, al Suwaidi Z, Nye PM, Parish T, Stoker NG. 1998. Inactivation of mycobacteriophage D29 using ferrous ammonium sulphate as a tool for the detection of viable Mycobacterium smegmatis and M. tuberculosis. Res Microbiol 149:487–95.

13. Oliveira IC, Almeida RC, Hofer E, Almeida PF. 2012. Bacteriophage amplification assay for detection of Listeria spp. using virucidal laser treatment. Braz J Microbiol 43:1128–36.

14. Xiong X, Zhang HM, Wu TT, Xu L, Gan YL, Jiang LS, Zhang L, Guo SL. 2014. Titer dynamic analysis of D29 within MTB-infected macrophages and effect on immune function of macrophages. Exp Lung Res 40:86–98.

15. Barr JJ. 2017. A bacteriophages journey through the human body. Immunol Rev 279:106–122.

16. Green SI, Gu Liu C, Yu X, Gibson S, Salmen W, Rajan A, Carter HE, Clark JR, Song X, Ramig RF, Trautner BW, Kaplan HB, Maresso AW. 2021. Targeting of Mammalian Glycans Enhances Phage Predation in the Gastrointestinal Tract. mBio 12.

17. Almeida G, Ravantti J, Grdzelishvili N, Kakabadze E, Bakuradze N, Javakhishvili E, Megremis S, Chanishvili N, Papadopoulos N, Sundberg LR. 2024. Relevance of the bacteriophage adherence to mucus model for Pseudomonas aeruginosa phages. Microbiol Spectr 12:e0352023.

18. Hill DB, Button B. 2012. Establishment of respiratory air-liquid interface cultures and their use in studying mucin production, secretion, and function. Methods Mol Biol 842:245–58.

19. Kesimer M, Kirkham S, Pickles RJ, Henderson AG, Alexis NE, Demaria G, Knight D, Thornton DJ, Sheehan JK. 2009. Tracheobronchial air-liquid interface cell culture: a model for innate mucosal defense of the upper airways? Am J Physiol Lung Cell Mol Physiol 296:L92–L100.

20. Hill DB, Vasquez PA, Mellnik J, McKinley SA, Vose A, Mu F, Henderson AG, Donaldson SH, Alexis NE, Boucher RC, Forest MG. 2014. A biophysical basis for mucus solids concentration as a candidate biomarker for airways disease. PLoS One 9:e87681.

21. Hill DB, Long RF, Kissner WJ, Atieh E, Garbarine IC, Markovetz MR, Fontana NC, Christy M, Habibpour M, Tarran R, Forest MG, Boucher RC, Button B. 2018. Pathological mucus and impaired mucus clearance in cystic fibrosis patients result from increased concentration, not altered pH. Eur Respir J 52.

22. Hancock LA, Hennessy CE, Solomon GM, Dobrinskikh E, Estrella A, Hara N, Hill DB, Kissner WJ, Markovetz MR, Grove Villalon DE, Voss ME, Tearney GJ, Carroll KS, Shi Y, Schwarz MI, Thelin WR, Rowe SM, Yang IV, Evans CM, Schwartz DA. 2018. Muc5b overexpression causes mucociliary dysfunction and enhances lung fibrosis in mice. Nat Commun 9:5363.

23. Ehre C, Rushton ZL, Wang B, Hothem LN, Morrison CB, Fontana NC, Markovetz MR, Delion MF, Kato T, Villalon D, Thelin WR, Esther CR, Jr., Hill DB, Grubb BR, Livraghi-Butrico A, Donaldson SH, Boucher RC. 2019. An Improved Inhaled Mucolytic to Treat Airway Muco-obstructive Diseases. Am J Respir Crit Care Med 199:171–180.

24. White A, Knight V. 1958. Effect of tween 80 and serum on the interaction of mycobacteriophage D-29 with certain mycobacterial species. Am Rev Tuberc 77:134–45.

25. Xu Z, Ding Z, Zhang Y, Liu X, Wang Q, Shao S, Liu Q. 2023. Shelf-life prediction and storage stability of Aeromonas bacteriophage vB_AsM_ZHF. Virus Res 323:198997.

