## Supplemental Methods for "Inactivating conditions of therapeutic mycobacteriophages"

Supplemental File

**Materials and Methods**

**Bacterial strains and media:**

Liquid cultures of *Mycobacterium smegmatis* mc^2^155 were grown at 37°C in Middlebrook 7H9 media (BD Difco™) supplemented with 0.5% glycerol, 0.2% glucose, and 0.05% Tween-80 (Fisher). Cultures were grown at 37°C with 80 rpm of agitation for 2 to 3 days prior to use in agar overlay assays to allow time for the *M. smegmatis* produced Tween hydrolase to degrade the Tween-80 (1). For growth on agar, Middlebrook 7H10 agar (BD Difco™) plates supplemented with 0.5% glycerol and 0.2% glucose were used.

**Phage stocks and high-titer phage lysate preparation:**

BPsΔ33HTH-HRM10 (BPsΔ), Muddy, and ZoeJΔ45 (ZoeJΔ) mycobacteriophages were used in the study (2). Phage propagation and quantification was performed with lawns of saturated stationary phase *M. smegmatis* mc^2^155 in top agar overlays that were prepared by mixing 0.3 ml of mc^2^155 culture and 100 µl of phage in 3.5 ml of molten 0.8% Noble top agar (ThermoScientific) poured on a 7H10 agar plate described above. Plates were incubated at 37°C and plaques counted 2 days later. To prepare high-titer phage lysates, phages were eluted from the agar overlay by flooding plates with mycobacteriophage (MP) buffer pH 7.5 (50 mM Tris/HCl pH 7.5, 150 mM NaCl, 10 mM MgSO_4_◦7H_2_O, 2 mM CaCl_2_) (1). The eluted phage was centrifuged at 3700 x g for 15 minutes to pellet debris and then filter sterilized by syringe passage through a 0.22 µm filter (Fisher; PVDF membrane). If necessary, the lysate was concentrated by one of two methods: ultracentrifugation or polyethylene glycol (PEG) precipitation. For ultracentrifugation, phage lysates were centrifuged at 175,000 x g at 4°C in an Optima XPN–80 ultracentrifuge with a Swinging-Bucket rotor (Beckman Coulter, Inc.). The phage pellet was resuspended in a smaller volume of MP buffer and rocked up to 3 days at 4°C before filtration through a 0.22 µm filter. For PEG precipitation, phages were concentrated as described previously (3). In brief, the phage lysate was precipitated overnight at 4°C with 4% polyethylene glycol (PEG) 8000 and 0.5 M NaCl and recovered by centrifugation at 3,700 x g for 30 min at 4°C. The precipitated phage was then resuspended in MP buffer overnight. Following a second spin to remove any undissolved material, the resuspended phage was transferred to an ultrafiltration concentrator with a 100 kDa MWCO (Pierce Protein Concentrator), washed with 40 mL of MP buffer, and concentrated to a smaller volume by centrifugation at 3,700 x g at 4°C. The concentrated phage ~10^9-10^ PFU/ml was recovered from the upper reservoir of the concentrator and filtered through a 0.22 µm filter.

**Sensitivity experiments:**

The following conditions were evaluated for their impact on phage titer at specific time points. With the exception of the temperature sensitivity experiments, treatments were performed at room temperature 22°C. At specific timepoints, treated and untreated phage samples were serially diluted in MP buffer in triplicate and spotted onto a solidified agar overlay with *M. smegmatis* to quantitate the level of viable phage +/- treatment.

Temperature. Phages in MP buffer were incubated in water baths at 22°C, 37°C, 45°C, 55°C, or 70°C.

Phage inactivation buffer (PIB) with different pH. Phage inactivation buffer (PIB) was prepared as a solution of 40 mM citric acid, 10 mM potassium chloride, and 135 mM NaCl, pH 3.0 (3) (4) (5). To prepare PIB at pH 4.0 or 5.0, the pH was adjusted with NaOH. For evaluating PIB, 10 µl of phage lysate was mixed with 90 µl of PIB to achieve a 1:10 dilution of phage in PIB.

MP buffer at acidic pH. Phage lysate was incubated in MP buffer adjusted to pH 2, 3, 4, or 5 with HCl.

Ethanol. For challenging phage sensitivity against ethanol, 10 µl of phage lysate was incubated in 90 µl of either 63% or 90% ethanol.

Reducing agents. For challenge with reducing agents, 10 µl of phage lysate was mixed with 90 µl 100 mM Tris(2-carboxyethyl)phosphine (TCEP, ThermoScientific) or 90 µl of 10 mM Dithiothreitol (DTT, ThermoScientific) to achieve final concentrations of 10 mM TCEP or 1 mM DTT. Unbuffered 10 mM TCEP at pH 1.9, or 10 mM TCEP buffered with NaOH to pH 7 was tested. Unbuffered 1 mM DTT T at pH 5.3, or DTT buffered with NaOH to pH 7.0 was tested.

Mucus. Phage lysate was incubated in 2%, 4%, and 6% mucus that was harvested from human bronchial epithelial (HBE) cells. Mucus was harvested from confluent HBE cultures by collecting PBS lavage and concentrating to 2, 4, and 6%(6).

Ferrous ammonium sulfate (FAS) and tannic acid (TA). A solution of 2.5 mM FAS (Sigma Aldrich) in 0.002% tannic acid (Sigma-Aldrich) was prepared along with separate solutions of 2.5 mM FAS or 0.002% tannic acid. 10 µl of phage lysate was added to 90 µl of 2.5 mM FAS or 0.002% tannic acid. 10 µl of phage lysate was also added to 90 µl of 2.5 mM FAS, 0.002% tannic acid solution for incubation. Final concentration of FAS was 2.25 mM.

Detergents. Equal volumes of phage lysate and 0.2% Tween-80 (ThermoScientific), 0.2% Tyloxapol (Sigma-Aldrich) or 0.2%, 2% or 20% Triton X-100 (Sigma-Aldrich) were mixed to achieve a final concentration of 0.1% - 10% for incubation.

Dimethyl sulfoxide (DMSO) (Sigma-Aldrich). Equal volumes of phage lysate and 2% DMSO solution prepared in MP buffer were mixed to achieve a final concentration of 1% DMSO.

Detergent addition to *M. smegmatis*. Liquid cultures of *M. smegmatis* were grown as described above to saturation, after which time the cells were pelleted at 3700 x g for 8 minutes and resuspended in equal volume of either Middlebrook 7H9 media supplemented with 0.5% glycerol, 0.2% glucose (no detergent), or the same Middlebrook 7H9 media with freshly added 0.1% Tween-80 media or 0.1% Tyloxapol. Agar overlays were prepared as above with each of the three different *M. smegmatis* culture preparations and 10-fold serial dilutions of each phage was spotted onto the agar overlays. Plates were incubated at 37°C. Images were taken after 22- and 46-hours incubation on the Bio-RAD GelDOC Go (software version 3.0.0.07) platform with the FAST Blast setting and the same exposure time of 0.09 seconds.

**Statistics and reproducibility**

Results were quantified as plaque forming units (PFU)/ml. The mean of three technical replicates for each time point was calculated and log transformed. Treatments that led to a significant reduction in level of phage compared to an untreated phage control at the same time point were determined by the log change (log PFU_treatment_ – log PFU_control_) for each of a minimum of two biological replicates. The limit of detection was 2,500 or 3.4 log PFU/ml. Statistical significance was determined by one-way analysis of variance (ANOVA, p<0.05) and the Tukey’s post-test for experiments with multiple timepoints. Experiments with a single timepoint were analyzed by unpaired t-test for comparing treated to untreated controls (p <0.05). GraphPad Prism 10 (version 10.5) was used for data analysis.

**References**

1.	Braunstein M, Bardarov SS, Jacobs WRJ. 2002. Genetic methods for deciphering virulence determinants of *Mycobacterium tuberculosis*, p 67-99. *In* Clark VL, Bavoil PM (ed), Methods in Enzymology, vol 358. Academic Press, London.

2.	Dedrick RM, Smith BE, Cristinziano M, Freeman KG, Jacobs-Sera D, Belessis Y, Whitney Brown A, Cohen KA, Davidson RM, van Duin D, Gainey A, Garcia CB, Robert George CR, Haidar G, Ip W, Iredell J, Khatami A, Little JS, Malmivaara K, McMullan BJ, Michalik DE, Moscatelli A, Nick JA, Tupayachi Ortiz MG, Polenakovik HM, Robinson PD, Skurnik M, Solomon DA, Soothill J, Spencer H, Wark P, Worth A, Schooley RT, Benson CA, Hatfull GF. 2022. Phage Therapy of Mycobacterium Infections: Compassionate-use of Phages in Twenty Patients with Drug-Resistant Mycobacterial Disease. Clin Infect Dis doi:10.1093/cid/ciac453.

3.	Schmalstig AA, Wiggins A, Badillo D, Wetzel KS, Hatfull GF, Braunstein M. 2024. Bacteriophage infection and killing of intracellular *Mycobacterium abscessus.* mBio 15:e0292423.

4.	Brindley MA, Maury W. 2008. Equine infectious anemia virus entry occurs through clathrin-mediated endocytosis. J Virol 82:1628-37.

5.	Zhang L, Sun L, Wei R, Gao Q, He T, Xu C, Liu X, Wang R. 2017. Intracellular *Staphylococcus aureus* Control by Virulent Bacteriophages within MAC-T Bovine Mammary Epithelial Cells. Antimicrob Agents Chemother 61.

6.	Hill DB, Button B. 2012. Establishment of respiratory air-liquid interface cultures and their use in studying mucin production, secretion, and function. Methods Mol Biol 842:245-58.
