## Supplemental Fig. 1 for "Inactivating conditions of therapeutic mycobacteriophages"

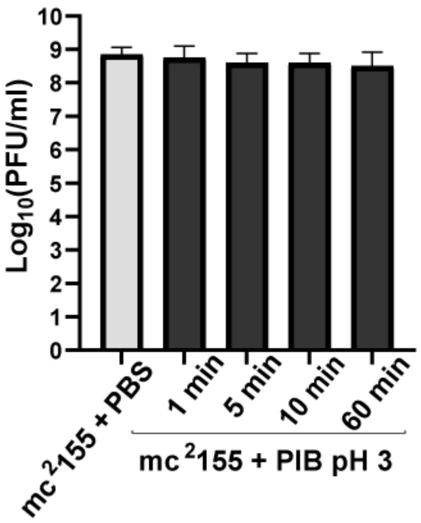


Supplemental Fig 1. *Mycobacterium smegmatis* is not sensitive to phage inhibition buffer (PIB) at pH 3. A saturated culture of *M. smegmatis* mc$^2$155 was pelleted, washed once in 1 X PBS, and resuspended in equal volume of 1 X PBS or in equal volume of PIB pH 3. The PIB pH 3 samples were incubated at room temperature for up to 60-minutes and colony forming units (CFU) were quantified. Results from two independent experiments (each with technical replicates) are plotted. No statistical significance was observed by one-way ANOVA ($p<0.05$) with Dunnett post-test.
